# The FSGS disease gene product and nuclear pore protein NUP205 regulates nuclear localization and activity of the transcriptional regulators YAP and TAZ in podocytes

**DOI:** 10.1101/2023.03.07.531564

**Authors:** Lioba Ester, Inês Cabrita, Michel Ventzke, Marita Christodoulou, Francesca Fabretti, Thomas Benzing, Sandra Habbig, Bernhard Schermer

## Abstract

**Background:** Mutations in genes encoding nuclear pore proteins (NUPs) cause steroid-resistant nephrotic syndrome (SRNS) and focal and segmental glomerulosclerosis (FSGS). The mechanisms of how NUP deficiency may cause podocyte dysfunction and failure of the kidney filtration barrier are elusive. The tightly controlled activity of the transcriptional effectors of the evolutionarily conserved Hippo pathway YAP and TAZ is essential for podocyte homeostasis. Here we analyze the role of NUPs in controlling YAP/TAZ nuclear import and activity in podocytes.

**Methods:** We used quantitative label-free mass spectrometry to characterize the YAP/TAZ interactomes in podocytes, particularly identifying NUP interactions. Moreover, we specifically studied NUP205 in controlling YAP/TAZ nuclear import and YAP/TAZ-dependent target gene expression.

**Results:** Here we identify the disease-causing nuclear pore proteins NUP107, NUP133, NUP205, and XPO5 as components of YAP and TAZ protein complexes in podocytes. We demonstrate that NUP205 is essential for YAP/TAZ nuclear import. The nuclear interaction of YAP/TAZ with TEAD1 and their transcriptional activity were dependent on NUP205 expression. Furthermore, we identify a feedback regulatory mechanism that controls YAP activity depending on TAZ-mediated NUP205 expression.

**Conclusion:** This study links the disease protein NUP205 with the activity of the transcriptional regulators and Hippo effectors YAP and TAZ and suggests a pathogenic role of YAP/TAZ-deregulation in podocytes in patients with *NUP205* mutations. Moreover, this study suggests an important role of YAP/TAZ signaling in human FSGS.

**SIGNIFICANCE STATEMENT:** Understanding the interference of signaling pathways in genetic diseases is essential to finally develop tailored treatment strategies. The increasing number of pathogenic variants causing focal-segmental glomerulosclerosis (FSGS) comprises genes encoding for proteins of the nuclear-cytoplamic shuttling machinery. We here found several of these proteins in an unbiased interactome analysis of the Hippo signaling effector proteins YAP and TAZ in podocytes, pointing to a connection between nucleoporins and the Hippo pathway. Further analyses confirmed that NUP205 is essential for correct YAP/TAZ shuttling; depletion of NUP205 resulted in cytoplasmic retention and inactivation of YAP/TAZ. We suggest the nuclear pore and hippo signaling as a pathogenic module in FSGS.

## INTRODUCTION

FSGS is one of the main histopathological findings in proteinuric renal diseases^1^. FSGS is caused by dysfunction and loss of podocytes disrupting the glomerular filtration barrier. Podocytes are terminally differentiated cells whose primary and secondary processes are essential for the formation of this barrier.^2, 3^ Cell-matrix adhesion firmly attaches these foot processes to the glomerular basement membrane.^4^ Adjacent podocytes are separated by filtration slits, bridged by membrane-like junctions called the slit diaphragm.^5^ Multiple causes were described to lead to foot process effacement and podocyte loss, including immunological and systemic processes and genetic variations.^6^ More than 50 mutations in various genes encoding for proteins of the slit diaphragm, cell membrane, cytoskeleton, and mitochondria have been identified to induce podocyte injury and the loss of integrity of the filtration barrier.^7, 8^ Recently, mutations in specific genes of the nuclear pore complex and the related shuttling machinery have been added to the list of genes causing steroid-resistant nephrotic syndrome (SRNS), a disease complex that includes the histopathological finding of FSGS.^9^

The nuclear pore complex (NPC) forms a channel at the nuclear membrane that allows nucleocytoplasmic transport in both directions.^10, 11^ The NPC consists of 30 different nuclear pore proteins called nucleoporins (NUPs).^12^ These NUPs interact in subclusters to form the major structural elements of the NPC: an inner, a nuclear and a cytoplasmic ring, the nuclear basket, and cytoplasmic filaments.^11^ In families suffering from SRNS, mutations in genes encoding for components of the inner ring (NUP205, NUP93), and of the cytoplasmic and nuclear rings (NUP85, NUP107, NUP133, NUP160) have been identified.^9, 13^ In addition, mutations in XPO5, one of the proteins involved in nuclear export acting in concert with nucleoporins, have been discovered.^9, 14^ However, the exact mechanism of how the disruption of the nucleocytoplasmic shuttling affects podocyte homeostasis remains to be elucidated. The Hippo Signaling pathway and its two effector proteins, Yes-associated protein (YAP1, hereafter referred to as YAP) and WW-domain-containing transcription regulator 1 (WWTR1, hereafter referred to by the alternative name TAZ), are important regulators of many cellular processes, such as cell differentiation, proliferation, and apoptosis.^15^ YAP and TAZ shuttle between the cytoplasm and the nucleus in response to various inputs. In the nucleus, YAP and TAZ act as transcriptional coactivators and regulate the expression of downstream targets in synergy with other transcription factors like TEA domain family members (TEAD1-4).^16^ In most healthy adult tissues, YAP and TAZ are usually maintained in an inactive state, i.e., phosphorylated and retained in the cytoplasm, only shuttling to the nucleus for regenerative or malignant growth.^17^ In podocytes YAP and TAZ seem to have divergent roles. Inputs that regulate YAP and TAZ activity, like cell-cell adhesions, cell polarity, or mechanical forces, are highly relevant in podocyte biology. Nuclear YAP activity has been shown to promote cell survival and inhibit podocyte apoptosis in FSGS.^18, 19^ Furthermore, mice lacking YAP or TAZ specifically in podocytes develop proteinuria and FSGS.^20, 21^ Therefore, podocyte homeostasis relies on a well-balanced nuclear shuttling and activity of YAP and TAZ, a process whose regulation is so far not well understood.^19, 22-27^

Here, we explore the interactome of YAP and TAZ in an *in vitro* mouse podocyte model. We identify several nuclear transport components that associate in protein complexes with both YAP and TAZ. Focusing on the FSGS protein NUP205, we describe its regulation of the nucleocytoplasmic shuttling of YAP and TAZ, relevant for their gene transcription activity. Lastly, we further identify a TAZ-indirect feedback regulation of YAP by NUP205. These results unmask a molecular pathomechanism perspective underlying FSGS.

## METHODS

### Cells

Heat sensitive mouse podocytes (hsMPs) were cultured in RPMI-1640 medium (Gibco, 61870) supplemented with 10% fetal bovine serum (FBS, Gibco, 10270), sodium pyruvate (Sigma, S8636), 20 mM HEPES (Sigma, H0887) and IFN-δ (0.25 µl/ml, PeproTech, 315-05). Podocytes were cultured on Collagen I-coated 10-cm primaria cell culture dishes (Corning, 353803), at low confluency and at 33°C, as previously described.^28^ The mouse podocyte cell line was obtained from Stuart Shankland (Seattle, WA). HEK293T cells were cultured in DMEM (Gibco, 31966) supplemented with 10% FBS at 37°C. For transfection experiments, cells were grown until 40% confluence and transfected with ON-TARGETplus SMARTpool siRNAs (Dharmacon) using Lipofectamine 3000 (Invitrogen, L3000-015). ON-TARGETplus Non-targeting pool (Dharmacon) was used as a negative control, scrambled RNA. The sequences of all siRNAs are described in Table 1. All experiments were performed 48h after transfection, unless mentioned otherwise. All cell lines were regularly tested for mycoplasma contamination by a regular kit. Cross-contamination with other cell lines was not observed and is currently not reported.

**Table 1.**
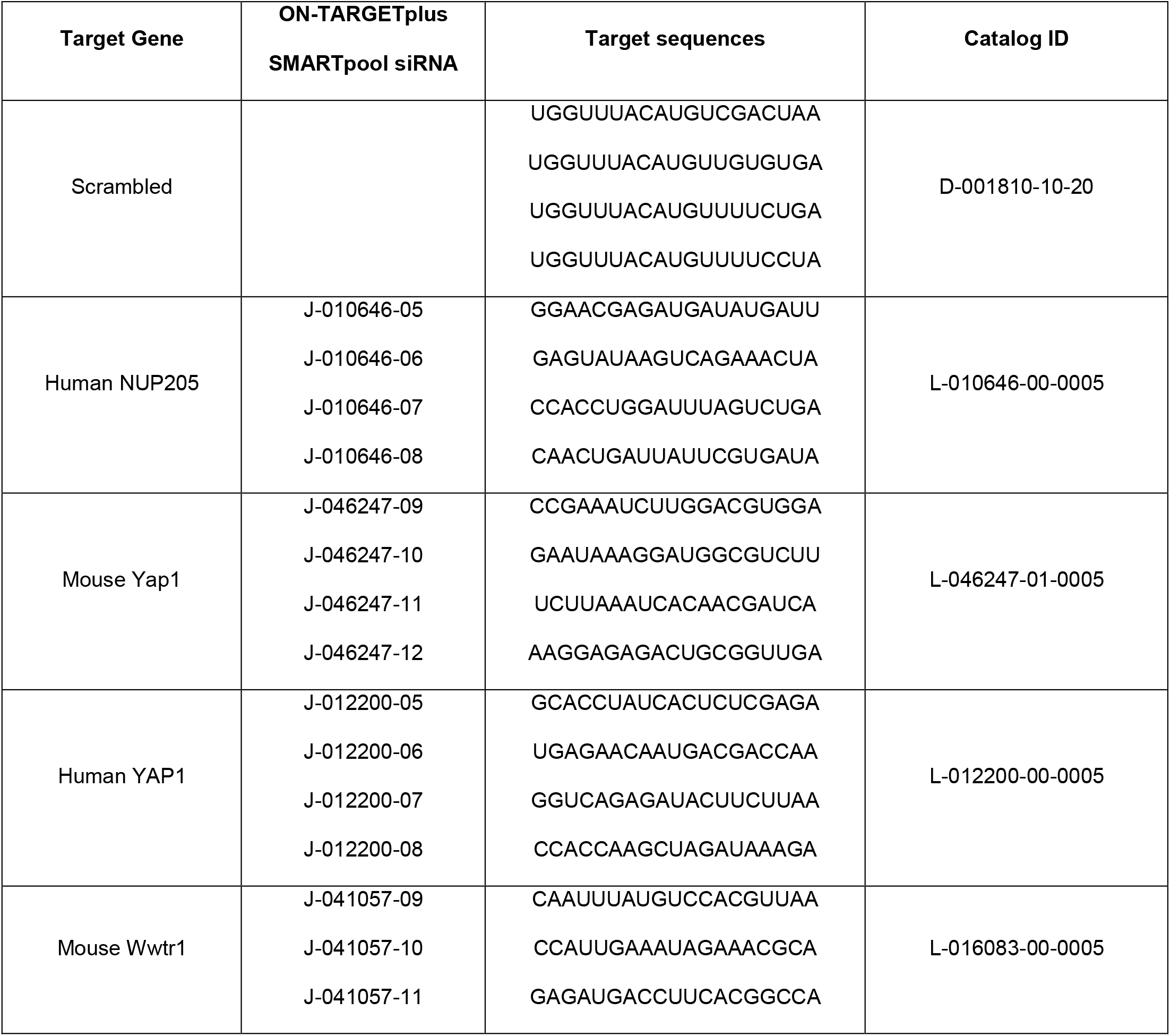
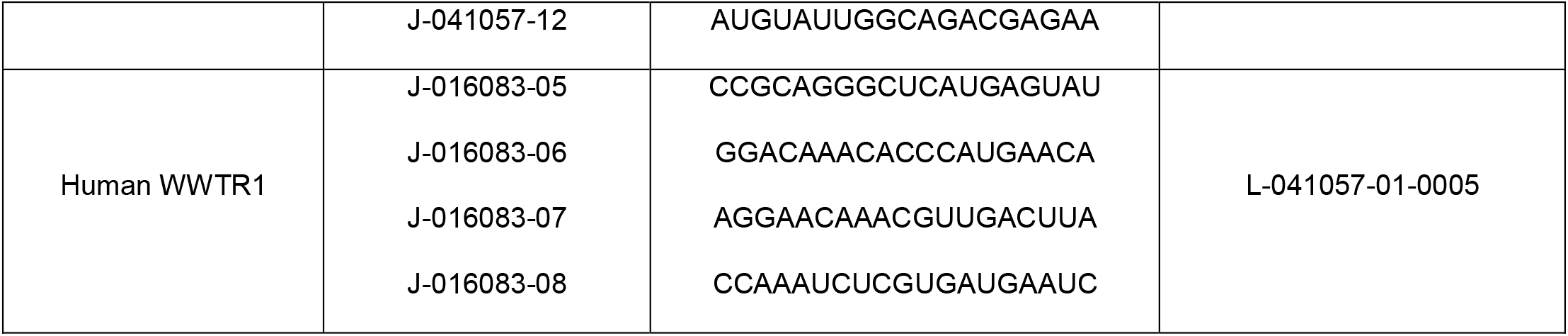
ON-TARGETplus SMARTpool siRNA sequences.

### Generation of stable cell lines

Using Transcription Activator-Like Effector Endonucleases (TALEN) technology, we generated stable hsMPs cell lines carrying a single copy integration of a transgenic construct expressing a flagged version of mouse Yap (3XFLAG.YAP), Taz (3XFLAG.TAZ) or Ruby (3XFLAG.RUBY) in the *Rosa26* locus. Rosa26_TV_CMV-3xFL-mRuby-STOP_EF1a-Puro-T2A-copGFP plasmid was based on pDonor MCS Rosa26 (gift from Charles Gersbach; addgene plasmid #37200).^29^ Four point mutations were added in the sequence of the later plasmid: HindIII site 2253 killed (now AATCTT), KpnI site 657 killed (now GGTAGC), NotI site 821 killed (now GGGGCCGC), NotI site 1492 killed (now GGGGCGGC). Mouse YAP or TAZ (kindly provided by Michael Yaffe) was inserted after excising of mRuby. The integration of the constructs in the mouse *Rosa26* locus was possible by co-transfecting with TALEN-mROSA26 KKR and TALEN-mROSA26 ELD (gift from Radislav Sedláček; addgene plasmid #60025 and #60026).^29, 30^ After transfection, stably transfected cells were selected with 1.5 µg/ml of puromycin (InvivoGen, ant-pr). Cells were maintained in the presence of puromycin and ready to be used. The correct expression of the tagged proteins was confirmed by immunoblotting (Supplemental Figure 1B).

**Figure 1.**
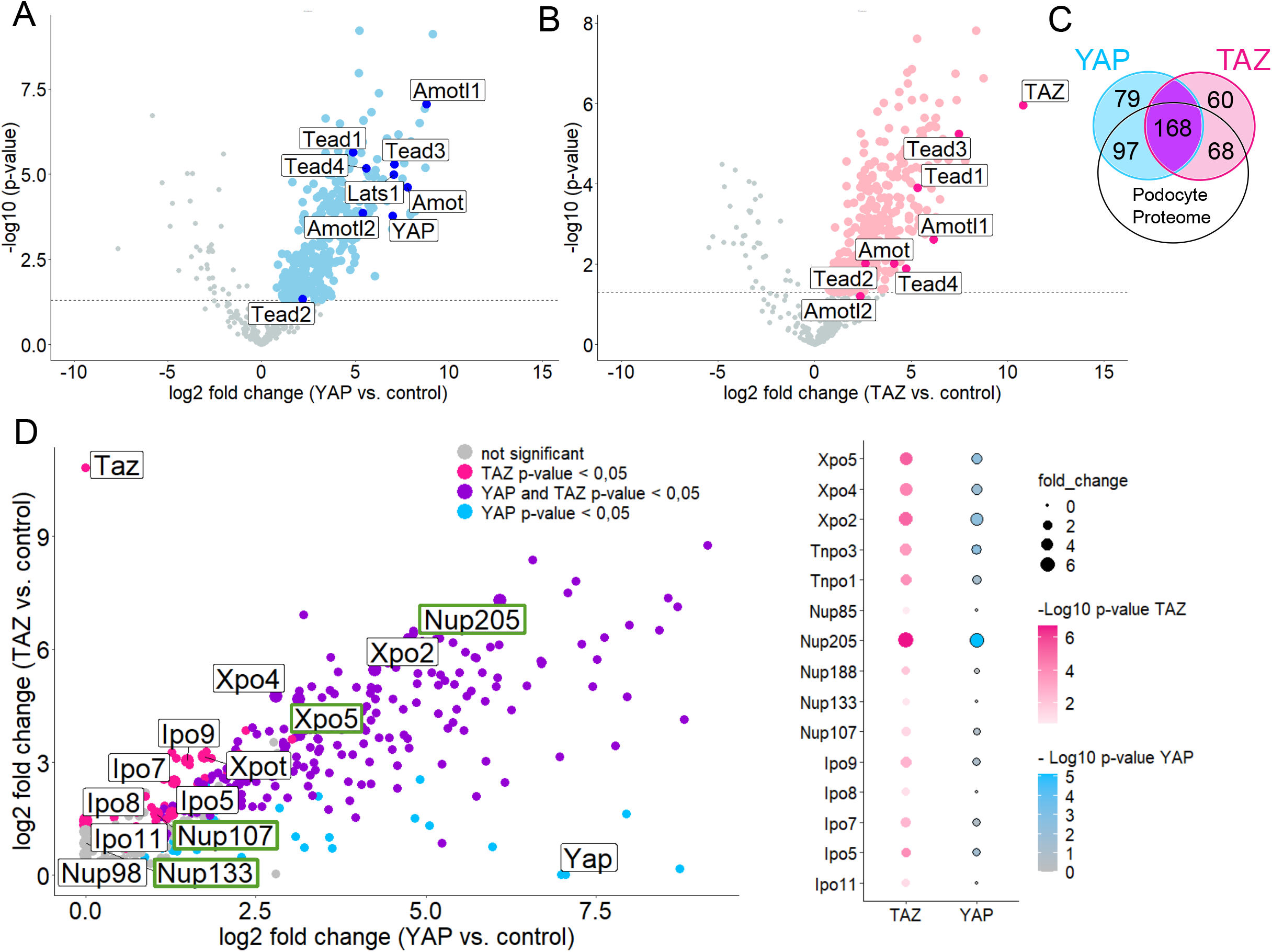
Podocyte interactome shows interplay of YAP and TAZ with nuclear transport proteins. Volcano plots showing the interactors of YAP (A) and TAZ (B) in immortalized podocytes (hsMPs) overexpressing 3xFLAG.YAP or 3xFLAG.TAZ. Logarithmized ratios are plotted against the negative logarithmic *p*-value of the Student’s *t*-test. Data points with a *p*-value<0.05 are printed bold (above dashed line). Known interactors are highlighted and labelled. (C) Venn diagram indicating numerical YAP (pink) and TAZ (blue) interactors overlapping with mouse podocyte proteome from Rinschen *et al*.^35^ (D) Scatter plot combining the interactors of YAP and TAZ, with shared interactors (*p*-value for YAP and TAZ < 0.05) highlighted in purple. Proteins of the nuclear-shuttling machinery are labelled and highlighted in green when related to FSGS. Dot plot showing proteins of nuclear shuttling machinery that are shared interactors by YAP and TAZ.

### Immunoprecipitation

For each replicate of the experiments, 2-4 ×10-cm dishes (60% confluency) were used. Cells were harvested with ice-cold PBS. The harvested cells were lysed in modified RIPA Buffer (1% Igepal NP40, 50 mM Tris·HCl pH 7.5, 150 mM NaCl, 0.25% Sodium Deoxycholate, and complete protease inhibitors without EDTA (PIM; Roche)). Cells were homogenized 8 times with a 27-gauge needle and incubated on ice for 15 min. Afterwards the samples were sonicated with a Bioruptor (10 min, cycle 30/30 sec) to degrade chromatin followed by centrifugation at 20,000xg for 30 minutes at 4°C and ultracentrifugation at 125,000xg for 30 minutes at 4°C. The samples for mass spectrometry (MS) analysis were incubated with Dynabeads protein G that were precoupled with the M2-Flag-antibody (Sigma, F1804) at 4°C overnight. Before adding of the antibody-coupled-beads a small aliquot of each sample was preserved and diluted with 2x SDS-PAGE sample buffer for immunoblot analysis. The next day, the beads were washed 5 times and then resuspended in 50µl of 5% SDS in PBS and boiled for 5 minutes. The magnetic beads were collected on a magnet. The eluate was further processed for MS analysis except a small aliquot that was used to validate the immunoprecipitation by immunoblotting. To prepare for MS analysis DTT was added to a final concentration of 5 mM, vortexed and incubated at 55°C for 30 minutes. After that CAA was added to a final concentration of 40 mM, vortexed and incubated in the dark for 30 minutes at room temperature. After centrifugation for 10 minutes at 20,000xg the supernatant was stored at -20°C to be then further processed and measured by the CECAD Proteomics Facility, University of Cologne. For YAP-immunoprecipitation, whole cell supernatants were incubated with 1 µg of rabbit IgG (Santa Cruz, sc-2027) or 0.2 µg of rabbit anti-YAP (Cell Signaling, 4912), overnight at 4°C. The next day Protein A sepharose beads were added and incubated for 1h at 4°C on an overhead shaker. Then, beads were washed 5 times and proteins eluted in 1x SDS-PAGE sample buffer for immunoblot analysis.

### Mass Spectrometry and data processing

Proteomics analysis were conducted at the CECAD Proteomics Facility, University of Cologne. All samples were analyzed on a Q-Exactive Plus (Thermo Scientific) mass spectrometer that was coupled to an EASY nLC 1000 or 1200 UPLC (Thermo Scientific). Samples were loaded onto an in-house packed analytical column (50 cm × 75 µm I.D., filled with 2.7 µm Poroshell EC120 C18, Agilent) that has been equilibrated in solvent A (0.1 % formic acid in water) and peptides were separated at a constant flow rate of 250 nL/min using a 50 min gradient followed by a 10 min wash with 95 % Solvent B (0.1% formic acid in 80% acetonitrile) for 10 minutes.

The mass spectrometer was operated in data-dependent acquisition mode. MS1 survey scans were acquired from 300 to 1750 m/z at a resolution of 70,000. The top 10 most abundant peptides were isolated within a 1.8 Th window and subjected to HCD fragmentation at a normalized collision energy of 27%. The AGC target was set to 5e5 charges, allowing a maximum injection time of 110 ms. Product ions were detected in the Orbitrap at a resolution of 35,000. Precursors were dynamically excluded for 10 seconds.

All mass spectrometric raw data were processed with Maxquant (version 1.5.3.8) using default parameters. Briefly, MS2 spectra were searched against the Mouse reference (downloaded at 16.6.2017), including a list of common contaminants. False discovery rates on protein and PSM level were estimated by the target-decoy approach to 1% (Protein FDR) and 1% (PSM FDR) respectively. The minimal peptide length was set to 7 amino acids and carbamidomethylation at cysteine residues was considered as a fixed modification. Oxidation (M) and Acetyl (Protein N-term) were included as variable modifications. The match-between runs option was enabled within replicate groups. LFQ quantification was enabled using default settings. Student’
ss T-tests were calculated in Perseus (version 1.6.1.1) after removal of decoys and potential contaminants. Data were filtered for at least 3 out of 3 values in at least one condition. Remaining missing values were imputed with random values from the lower end of the intensity distribution using Perseus defaults.

### Cellular fractions

For whole cell lysates, hsMPs or HEK293T cells were harvested with ice-cold PBS and lysed in modified RIPA Buffer. Cells were titurated 8 times with a 27-gauge needle and incubated on ice for 15 minutes. After centrifugation at 20,000xg for 15 minutes at 4°C the supernatant was mixed with 2x SDS-PAGE sample buffer and boiled for 5 minutes at 95°C.

Nuclear and cytoplasmic fractions were collected as previously described.^31^ Briefly, HEK293T cells were harvested on ice, after siRNA transfection in 7 ml ice-cold PBS. 1 ml of the resultant cell suspension was collected for processing as a whole cell lysate in SDS-PAGE sample buffer. The remaining cell suspension was pelleted at 48xg 5 minutes at 4°C. The supernatant was discarded, and the cell pellet was resuspended in 150 μl of the hypotonic Cell Fraction Buffer (10 mM HEPES, 1.5 mM MgCl_2_,10 mM KCl, pH 7.9 plus PIM (Roche)). Cells were incubated on ice for 10 minutes, and then disrupted 14 times using a 21-gauge needle. The mixture was gently centrifuged at 100xg for 30 minutes at 4°C to pellet the nuclear fraction. The supernatant was ultracentrifuged at 100,000xg for 30 minutes at 4°C. 80 μl of the resultant supernatant was transferred to a new 1.5 Eppendorf tube and mixed with 80 μl of 2×SDS-PAGE sample buffer, boiled for 5 minutes at 95°C, and centrifuged for 1 minute at 20,000xg to yield the cytosolic fraction. Meanwhile, the nuclear pellet was resuspended in 1 ml cold PBS and centrifuged at 9200xg for 10 minutes at 4°C. The supernatant was discarded, and the remaining pellet was washed once with PBS and centrifuged again for 10 minutes 9200xg at 4 °C. The pellet was then resuspended in 100 μl of 2x SDS-PAGE sample buffer, boiled for 10 minutes at 95°C and centrifuged for 5 minutes at 20,000xg to yield the nuclear fraction. 20 μl of the supernatant was analysed by Western Blot.

### Immunoblotting

Proteins were separated by 8.5% Sodium dodecyl sulphate (SDS) polyacrylamide gel electrophoresis and transferred onto a polyvinylidene fluoride membrane (Millipore, Immobilon-P, IPVH00010) using a semi-dry transfer unit (PEQLAB). After blocking in 5% bovine serum albumin, membranes were stained with 1:2000 of primary anti-beta-tubulin mouse monoclonal antibody (DSHB, E7), anti-fibrillarin rabbit monoclonal antibody (Abcam, ab166630), anti-NUP205 rabbit polyclonal antibody (Biomol, A303-935A), anti-YAP rabbit polyclonal antibody (Cell Signaling, 4912), anti-TAZ rabbit monoclonal antibody (Cell Signaling, 72804), anti-TEAD1 mouse monoclonal antibody (BD Biosciences, 610922), anti-beta-actin mouse monoclonal antibody (Cell Signaling, 3700 S) or anti-GAPDH rabbit monoclonal antibody (Cell Signaling, 5174), overnight at 4°C. Proteins were visualized using horseradish peroxidase-conjugated secondary antibody (1:30000) and ECL detection. Images were acquired with a Fusion Solo S (Vilber Lourmat Germany GmbH, Eberhardzell, Germany). Densitometric analysis was performed using ImageJ/Fiji Software version 2.9.0/1.53t (NIH, Bethesda, MD).^32^

### qPCR

RNA was isolated using TRIzol (Sigma, S8636) and the Direct-zol RNA MiniPrep (ZYMO research, R2052) according to the manufacturer’s instructions. In-column digestion with DNAse was performed for all samples to remove genomic DNA contamination. Quality and quantity of eluted RNA was checked using a NanoDrop spectrophotometer (PEQLAB). Complementary DNA (cDNA) was synthesized using the same amounts of RNA using the High-capacity cDNA reverse transcription (Thermo Fisher Scientific, 4368814). qPCR was performed using SYBR Green assays (Thermo Fisher Scientific, 4367659) and analyzed using a QuantStudio 12K Flex cycler system. Primer sequences are listed in Table 2. Quantification of relative expression levels was performed using the 2^−ΔΔCT^ method as previously described^33^, which involves normalization to a housekeeping gene. The gene *Hprt* was used as housekeeping gene. Afterward, expression was normalized to control condition, resulting in the final “relative mRNA expression” parameter. Primer pairs were tested for efficiency.

**Table 2.**
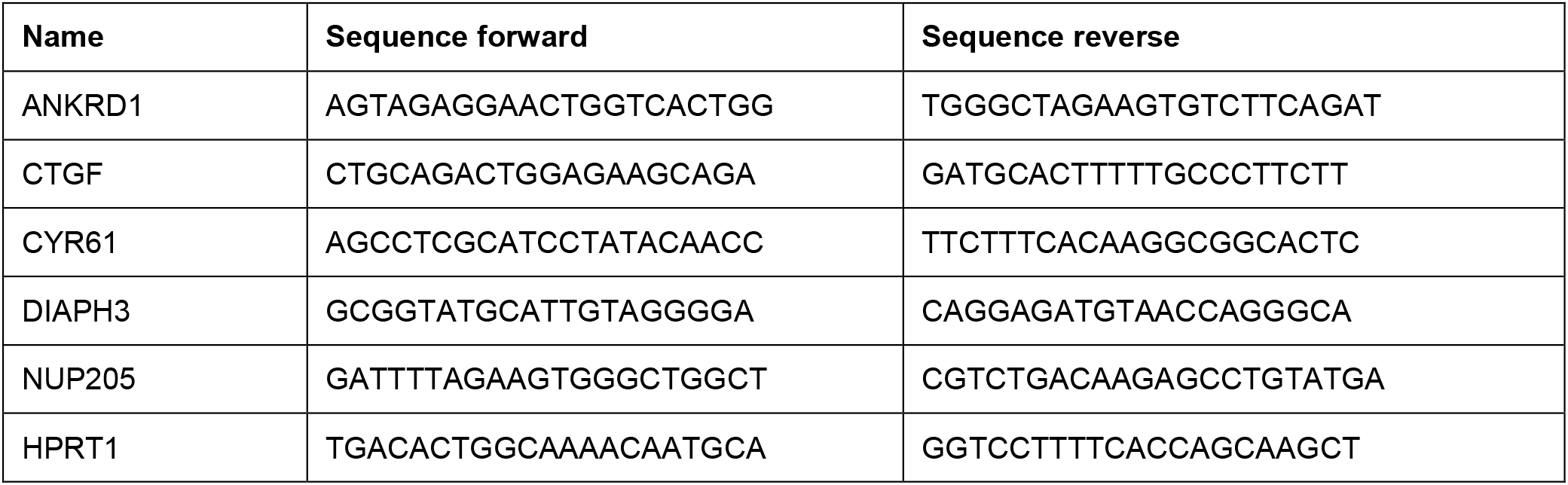
Primers used for qPCR.

### Immunocytochemistry

HEK293T cells were seeded onto 12 mm coverslips and transfected using indicated siRNA. 16h or 48h afterward, cells were rinsed with PBS and fixed with 4% formalin for 15 min. After washing with PBS, cells were permeabilized in 0.5% Triton X-100 in PBS, for 10 min. After blocking with 1% normal donkey serum (NDS, Biozol, END9010) and 1% bovine serum albumin (BSA, Thermo Fisher Scientific, 23209) in PBS, cells were sequentially co-stained with 1:200 primary anti-NUP205 rabbit polyclonal antibody (Biomol, A303-935A), anti-YAP mouse monoclonal antibody (Santa Cruz, sc-376830) or anti-TAZ mouse monoclonal antibody (BD Biosciences, 560235). Afterward, the coverslips were mounted with Prolong gold with DAPI (Invitrogen) and subjected to immunofluorescence microscopy. Pictures were taken with a STELARRIS 5 LIAchroic inverse confocal microscope (Leica) and objective HC PL APO 63x/1.30 GLYC CORR CS2 was used for acquisition. Subsequent image processing and analysis was performed using ImageJ/Fiji Software version 2.9.0/1.53t (NIH, Bethesda, MD).^32^ Nuclear and cytosolic expression levels were quantified by analysing fluorescence intensities in the regions of interest (ROI) and were plotted as proportions of nuclear versus cytoplasmic fluorescence.

### Statistical analysis

Data are reported as mean ± SEM. Student’s *t*-test for unpaired samples was used for statistical analysis. p < 0.05 was accepted as significant difference.

### Data availability

The mass spectrometry proteomics data have been deposited to the ProteomeX-change Consortium via the PRIDE partner repository with the data set identifier PXD040448.^34^

## RESULTS

### YAP and TAZ interact with the nuclear shuttling machinery in podocytes

To unravel the importance of YAP and TAZ in podocytes, we aimed to identify novel podocyte-specific components of the YAP/TAZ protein complexes. Therefore, we used a single-copy integration systems targeting the *Rosa26* locus and generated podocyte cell lines stably expressing low levels of 3xFLAG.YAP, 3xFLAG.TAZ or 3xFLAG.Ruby as negative control (Supplemental Figure 1A). Pulldown assays with Flag-antibody successfully immunoprecipitated FLAG-tagged Yap, Taz or Ruby as confirmed by immunoblotting (Supplemental Figure 1B). Five biological replicates generated on different days were analyzed together by mass spectrometry to identify specific components of Yap and Taz protein complexes in podocytes. Principal component analysis (PCA) revealed that all samples within a group (i.e., RUBY, YAP and TAZ) clustered together while the groups were markedly separated (Supplemental Figure 1C). Further analysis led to the identification of a large number of proteins enriched in either the YAP or TAZ dataset as compared to the control (Figure 1A/B). As expected, Yap and Taz were among the most prominently enriched proteins in the 3XFLAG.YAP pull-down (Figure 1A) and in the 3XFLAG.TAZ pull-down (Figure 1B), respectively. Taken together, our datasets revealed 362 proteins significantly enriched in the Yap-samples and 314 proteins in the Taz-samples as novel components of YAP/TAZ protein complexes. Among these identified proteins many well established interactors of Yap and Taz were included, like proteins of the Tead family, different angiomotins or large tumor suppressor kinase 1 (Lats1) verifying the accuracy of the data sets (Figure 1A/B). Furthermore, comparison of our data set with a published proteome expression profile of mouse podocytes isolated directly from glomeruli confirmed the expression of the majority of the identified proteins from our study in podocytes *in vivo*, underlining the relevance of our datasets for podocyte biology (Figure 1C). ^35^ Notably, the identified protein complexes associated with YAP and TAZ showed similarities which are well explained by their homology and common functions. Strikingly, among the shared proteins were numerous proteins of the nuclear shuttling machinery, like importins (Ipo), exportins (Xpo), transportins (Tnpo) and nucleoporins (Nup) that form the nuclear pore complex (NPC) (Figure 1D). Among those were NUP107, NUP133, NUP205 and XPO5. Variants in the corresponding genes have recently been identified to cause FSGS, making them even more interesting for podocyte biology.^9, 13, 36^ Specifically, Nup205 is the most abundant nuclear transport protein within the complex of both, Yap and Taz.

### NUP205 regulates YAP and TAZ nuclear shuttling

NUP205 is a protein from the inner ring subunit of the NPC, important for its assembly and function in active bidirectional transport of molecules between the nucleus and cytoplasm^37, 38^ To investigate the influence of this specific nucleoporin on the nuclear shuttling of YAP and TAZ we targeted endogenous NUP205 using siRNA and analyzed changes in endogenous YAP or TAZ subcellular localization. Subcellular fractionation assays revealed that knockdown of NUP205 reduced the nuclear expression of YAP and TAZ (Figure 2A). Meanwhile, the cytoplasmic expression levels of YAP and TAZ were increased, while total protein levels remained equal (Supplemental Figure 2A/B). The lack of constitutive nuclear shuttling of YAP and TAZ upon knockdown of NUP205 was further confirmed by assessing the shift in localization of endogenous proteins by immunofluorescence (Figure 2B). From these results, it is evident that NUP205 is essential for the nuclear import of both YAP and TAZ.

**Figure 2.**
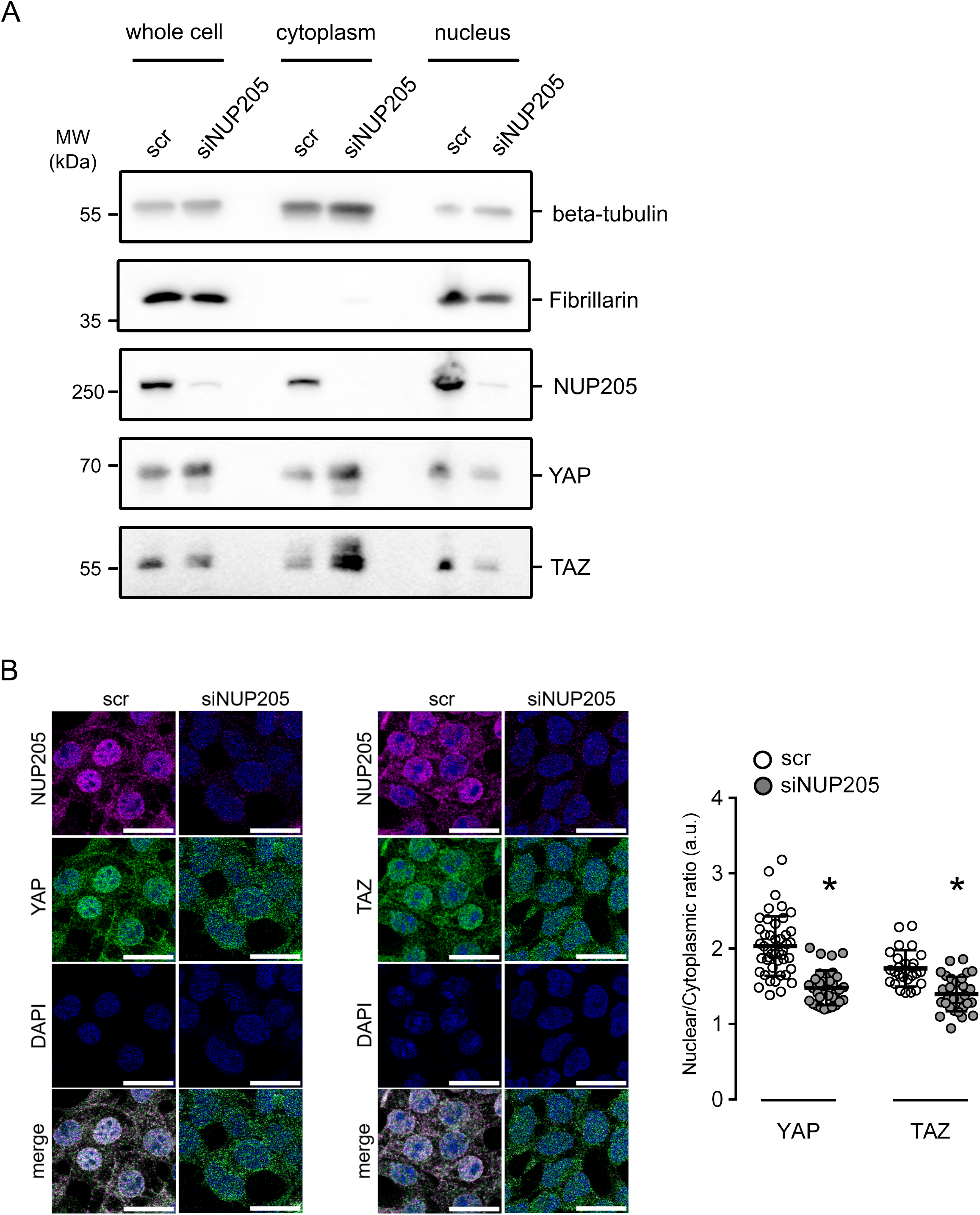
Downregulation of NUP205 compromises nuclear localization of YAP and TAZ. (A) Immunoblot showing the results of the cell fractionation assay for HEK293T cells transfected with siRNA NUP205 compared to control. The presence of the cytoplasmic protein beta-tubulin and the nuclear protein fibrillarin in the appropriate fraction was confirmed by western blotting with the corresponding antibodies. Downregulation of NUP205 decreases nuclear shuttling of YAP and TAZ, which stay confined in the cytoplasm. (B) Immunofluorescence staining of HEK293T cells transfected with siRNA NUP205 shows a decrease of nuclear YAP and TAZ compared to control. Co-staining in magenta for endogenous NUP205, in green for endogenous YAP (left) or TAZ (right), and nuclear stain with Hoechst (blue). All scale bars=20 µm. Summary of nuclear/cytoplasmatic fluorescence ratio quantification (n=26-45 cells). Data are reported as mean ± SEM. Representative data from three independent experiments are shown. *Significant decrease of nuclear staining of YAP and TAZ (unpaired *t*-test; *p*<0.05).

### Lack of NUP205 leads to a decrease in YAP and TAZ-dependent transcriptional activity

YAP and TAZ shuttle into the nucleus to function as co-transcription factors and initiate the transcription of various target genes.^39-41^ YAP/TAZ can contact the DNA only indirectly through transcription factor partners. One pivotal transcription factor they bind is TEAD1.^16^ TEAD1 is localized to the nucleoplasm, therefore, YAP needs to shuttle into the nucleus. To determine whether NUP205 downregulation causes a reduced YAP/TEAD1 nuclear interaction, TEAD1 was co-immunoprecipitated with YAP after NUP205 knockdown and control cells. Remarkably, YAP pulled down TEAD1 in control cells, and co-immunoprecipitation was diminished upon NUP205 knockdown (Figure 3A). In support of this, we found that the RNA levels of the bona fide target genes of YAP and TAZ – CYR61, DIAPH3, CTGF, and ANKRD1 - were reduced upon knockdown of NUP205 (Figure 3B). Taken together, depletion of NUP205 led to a reduction of nuclear YAP and TAZ and, consequently, a reduced interaction with TEAD1 and inhibition of target genes expression, confirming that the transcriptional activity of YAP and TAZ is dependent on the proper expression of the nuclear pore component NUP205.

**Figure 3.**
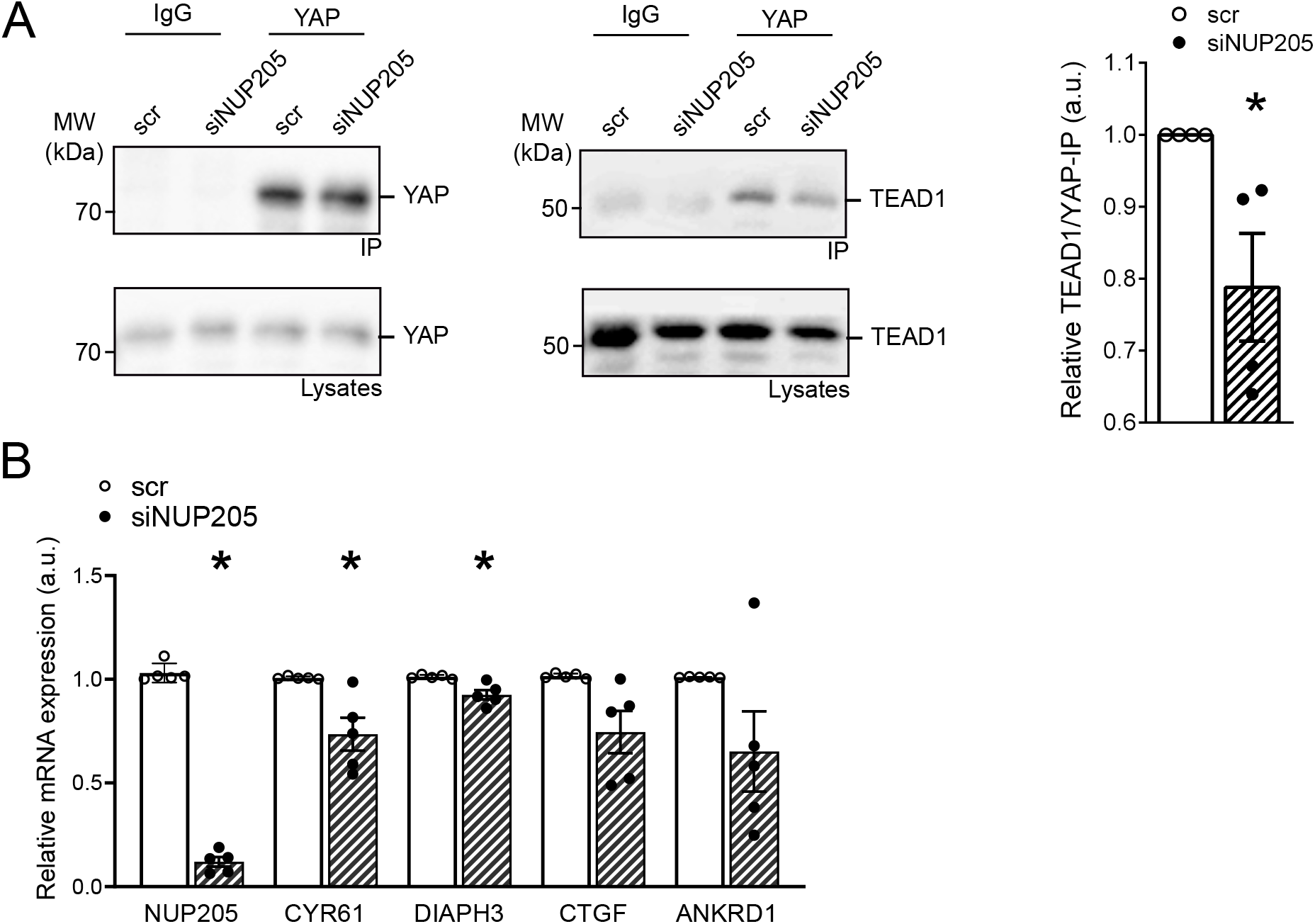
YAP and TAZ activity is decreased upon NUP205 downregulation. (A) Co-immunoprecipitation of endogenous YAP shows a decrease in interaction with TEAD1 in HEK293T cells transfected with siRNA NUP205 compared to control. IgG is used as negative control for the IP. Quantification of relative TEAD1 pulldown normalized to strength of YAP-immunoprecipitation (n=4). (B) qPCR shows decrease of relative mRNA expression of YAP/TAZ target genes in HEK293T cells transfected with siRNA NUP205 compared to control (n=5). Data are reported as mean ± SEM. *Significant decrease by siRNA NUP205 (unpaired *t*-test; *p*<0.05).

### Downregulation of TAZ impairs NUP205-dependent nuclear shuttling of YAP

Protein complex partners can mutually affect expression levels, stability, and function. Therefore, we further analyzed the effect of YAP or TAZ knockdown on NUP205. In cultured podocytes, knockdown of Taz, but not Yap, resulted in a significant reduction of Nup205 expression after 48 h (Figure 4A). To characterize the time course of this NUP205 reduction, we compared different instances of TAZ knockdown in HEK293T cells. Firstly, the immunoblotting revealed the chronological sequence of TAZ depletion upon transfection with TAZ siRNA (Figure 4B). While after 6 h of knockdown TAZ protein levels were still equivalent to control, a strong reduction could be observed from 12 h on. A global depletion of TAZ was obtained after 24 h. The decrease of NUP205 expression levels followed TAZ downregulation with a stronger transient reduction after 16 h. To examine whether downregulation of TAZ indirectly affected YAP subcellular localization through NUP205 downregulation, we again performed subcellular fractionation assays. We selected a period of 16 h for the TAZ knockdown because it resulted in the most efficient reduction of NUP205 expression. After 16 h of siRNA, not only TAZ, but also NUP205 showed a reduction in protein expression in the whole cell samples compared with control (Figure 4 C). Strikingly, NUP205 also showed a shift in subcellular localization from the nuclear to the cytoplasmatic fractions. YAP whole cell expression levels were unaffected by the knockdown of TAZ, but it was retained in the cytoplasm, showing an increased expression in the cytoplasmic fraction while being reduced in the nucleus. This confirmed that TAZ was indispensable for proper NUP205 expression and localization, which in turn affected the nuclear shuttling of YAP. To validate this effect, we examined the same conditions by immunofluorescence staining (Figure 4D). Once again, downregulation of TAZ affected NUP205 expression levels, quantified by whole cell fluorescence intensity. Notably, cells showing a reduced NUP205 expression also displayed a reduced nuclear expression of YAP, confirmed by assessing the nuclear/cytoplasmatic ratio. Taken together, the present results confirm the two-step regulation of YAP localization by TAZ through NUP205, the NPC protein essential for the shuttling of the Hippo pathway effector proteins.

**Figure 4.**
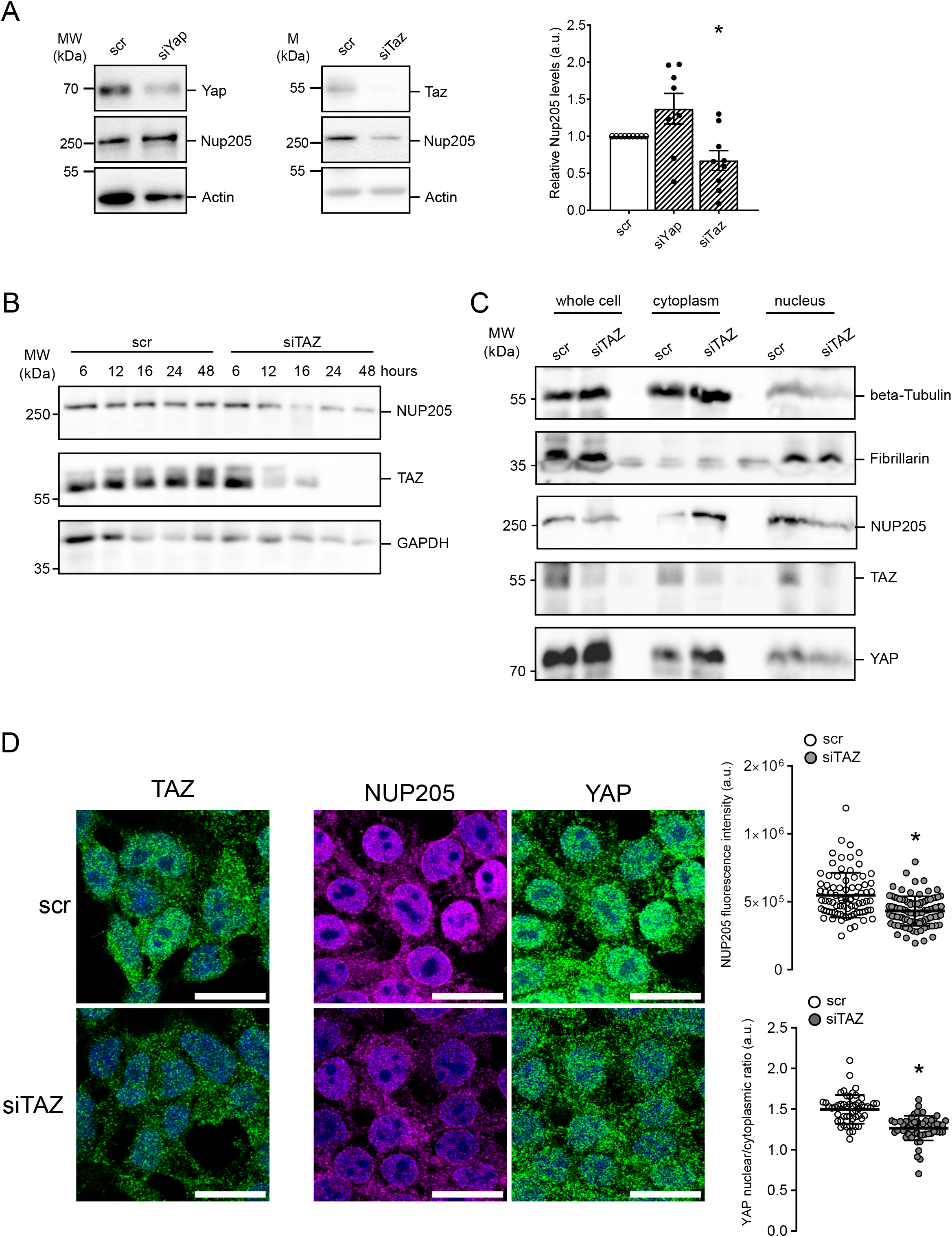
Lack of TAZ disturbs NUP205 expression and shifts YAP localization. (A) Immunoblot showing whole cell expression levels of endogenous NUP205 in hsMPs transfected with siRNA YAP or TAZ compared to control for 48h. Densitometry analysis of NUP205 expression normalized to actin (n=8-9). Data are reported as mean ± SEM. *Significant decrease by siRNA TAZ (unpaired *t*-test; *p*<0.05). (B) Immunoblot showing endogenous expression levels of NUP205, TAZ, YAP and GAPDH, in HEK293T cells transfected with siRNA TAZ compared to control, after 6, 12, 16 and 24h. Representative blot for n=3. (C) Immunoblot showing downregulation of NUP205 and the nuclear-to-cytoplasmatic-shift of YAP upon downregulation of TAZ after 16h in HEK293T cell fractionation assay. Representative blot for n=3. (D) Immunofluorescence staining of HEK293T cells transfected with siRNA TAZ for 16h compared to control. TAZ downregulation on the left. Co-staining in magenta for endogenous NUP205, in green for endogenous YAP, and nuclear stain with Hoechst (blue). Representative data from three independent experiments are shown. All scale bars=20 µm. Summary of Correlated Total Cell Fluorescence (CTCF) of NUP205 (n=81-95 cells) and summary of YAP nuclear/cytoplasmatic ratio quantification (n=53-57 cells). Data are reported as mean ± SEM. *Significant decrease by siRNA TAZ (unpaired *t*-test; *p*<0.05).

## DISCUSSION

Mutations in genes encoding for components of the NPC have been identified to cause FSGS, while the exact role of the nuclear-shuttling machinery in podocytes remains elusive.^9, 13, 36, 42^ NPCs regulate the transport of molecules shuttling from the cytoplasm to the nucleus, but have also been shown to be involved in transcriptional regulation.^43^ By exploring our podocyte interactome data of YAP and TAZ we identify novel associations of NUPs and nuclear transport receptors (NTRs), like importins and exportins, with YAP and TAZ. This is captivating because YAP and TAZ homeostasis, in addition to phosphorylation, depends on smooth nucleocytoplasmic shuttling.^15^ So far, not much is known on this direct regulation of the nuclear translocation of YAP and TAZ besides a dependency on NUP37 in hepatocellular carcinoma cells and oocytes, and a reported regulation of the nuclear pore size followed by the entry of YAP by mechanical forces.^44-46^ Further, Ipo7, which also was identified in our podocyte interactome data, has been shown to regulate the import of YAP in retinal pigment epithelium (RPE) cells.^47^ Among the newly identified nuclear-shuttling components of the YAP/TAZ complexes in our data are various proteins, the mutations of which are known to cause FSGS, like XPO5, NUP107 and NUP166, and NUP205.^9, 13^ Of those, NUP205 showed the most robust enrichment in both protein complexes. Four different mutations of NUP205 have been described so far to cause FSGS. ^7, 9, 48, 49^ Braun *et al*. were the first to identify NUP205 and at the same time NUP93 to cause FSGS and they suggested a role of NUP93 in the resistance to oxidative stress and the nuclear shuttling of SMAD.^9^ The consequences of NUP205 mutations on nuclear shuttling in podocytes, despite a reduced interaction with NUP93, had not been studied. Transport of proteins through the NPC is regulated by NUPs which are rich in phenylalanine-glycin repeats (FG-NUPs). FG-NUPs form liquid-liquid phase separates at the inner channel of the pore and impair free diffusion of macromolecules.^50^ They interact with NTRs to allow them and their cargo proteins to translocate rapidly in an energy-dependent manner.^51^ Theoretically, YAP and TAZ should, based on their size (55kDa), be imported into the nucleus assisted by NTRs. However, YAP and TAZ lack a classic nuclear localization signal that is thought to be crucial for nuclear translocation coupled to NTRs.^52^ Interestingly, NUP205 can transport macromolecules independently of NTRs. NUP205 has been reported to interact directly with the arginine-rich motif (ARM) of adenovirus for its nuclear import.^53^ Moreover, Nup192 (NUP205 homolog in yeast) has been shown to carry structural and functional similarities with NTRs and can thus translocate proteins through intact NPCs in an NTR-independent way by facilitated diffusion.^54^ We postulate that NUP205 forms a protein complex with YAP and TAZ, allowing their nuclear import through translocation of NUP205 within the pore as a dominant shuttling process in healthy podocytes (Figure 5A).

**Figure 5.**
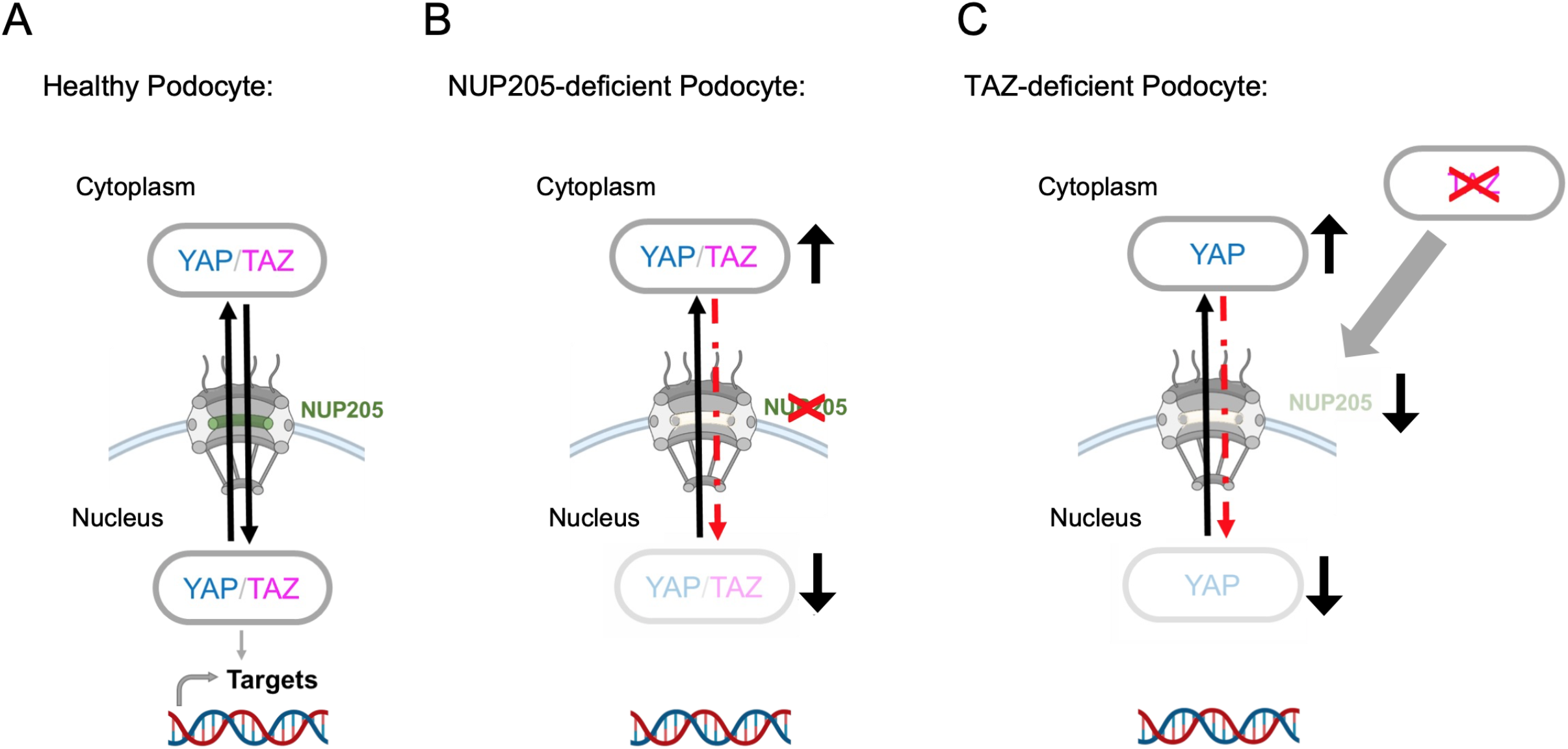
NUP205 is critical for nuclear shuttling of YAP in healthy podocytes. (A) In a healthy podocyte NUP205 is located at the nuclear membrane as part of the nuclear pore complex where YAP and TAZ can shuttle between cytoplasm and nucleus. (B) In the diseased podocyte lacking proper NUP205 expression, the nuclear translocation of YAP and TAZ is disrupted. YAP and TAZ are “trapped” in the cytoplasm resulting in a reduced nuclear transcriptional activity. (C) TAZ expression is crucial for NUP205 expression in podocytes. Reduced TAZ expression leads to reduced NUP205 expression, which then results in a shift of YAP localization. Created with BioRender.com.

For podocyte homeostasis, nuclear localization and activity of YAP have been reported to be essential for protecting the podocyte from apoptosis.^18, 25^ The importance of balanced YAP activity in podocytes is further supported by studies showing an initial increase of YAP activity followed by a decrease in rodent FSGS disease models.^19, 22, 27^ This temporary upregulation before the onset of disease suggests a transient protective role of YAP in the podocyte. Furthermore, increased activity of the Hippo Signaling cascade, that leads to an increased phosphorylation and cytosolic retention of YAP, consequently decreased its activity, led to disruption of the cytoskeletal integrity and podocyte morphology.^55^ Overall, YAP and TAZ seem to have a protective, antiapoptotic role, and their nuclear localization is pivotal for the healthy podocyte. Here, we show that NUP205 is not only required for the nuclear import of YAP and TAZ but is also crucial for their transcriptional activity in cooperation with TEAD1. Our results present a potential FSGS pathomechanism: when NUP205 fails to regulate the indispensable nuclear shuttling and activity of YAP and TAZ, it results in loss of protective properties thus, podocytes injury (Figure 5B).

Our identification of the association of NUPs with YAP and TAZ in podocytes led us to also investigate if YAP and TAZ influence the regulation of the nuclear import and export in podocytes. Recently it was shown that YAP regulates Ipo7, a NTR, by monopolizing its nuclear import in response of mechanical cues in RPE cells.^47^ To our knowledge, our data describes for the first time the regulation of NUPs by components of the Hippo Signaling pathway. We demonstrate that NUP205 expression is dependent on TAZ in podocytes. Moreover, our findings reveal an indirect decrease of YAP nuclear translocation by TAZ, suggesting a negative feedback regulation (Figure 5C).

Intriguingly, mutations in specific nucleoporins can lead to a very distinct cell-specific phenotype.^11^ Even more striking, different mutations in the same gene can cause fundamentally different phenotypes. Some NUP205 mutations lead to developmental disorders like abnormal cardiac left-right patterning, some contribute to a variety of neurological diseases,^56-59^ while others lead to SRNS.^13, 48, 60^ This suggest that nucleoporins have explicit roles in specific cell types and that certain cell types react peculiarly to changes in signaling of specific pathways. Strikingly, all those conditions caused by NUP205 mutations are related with YAP/TAZ expression.^61-68^

In conclusion, our findings highlight the important role of YAP/TAZ signaling in podocyte homeostasis by connecting the shuttling of YAP and TAZ to a nucleoporin, known to cause FSGS. Our work identifies a novel interdependency between transcription factors and NPCs which highlights new specific therapeutic targets for FSGS like stabilizing nuclear YAP activity in podocytes.

## AUTHOR CONTRIBUTIONS

L.E., I.C., B.S. S.H. and T.B. designed the study; L.E, I.C., F.F., M.C. and M.V. carried out experiments; L.E., I.C. analyzed the data; L.E. made the figures; L.E. and I.C. drafted the paper; L.E., I.C., T.B., S.H. and B.S. revised and approved the final version of the manuscript.

## ACKNOWLEDGMENTS

We would like to thank Angelika Köser for expert technical assistance. We acknowledge the help of the CECAD imaging facility and the CECAD proteomics facility. In addition, we would like to express our gratitude to all members of our laboratory for helpful discussions and support.

## DISCLOSURES

All authors declared no competing interests.

## FUNDING

This work was supported by the German Research Foundation (DFG) in the framework of the clinical research unit 329 (KFO 329; RP7 to S.H. / B.S and RP1 to T.B.). L.E. was supported by the Koeln Fortune program/Faculty of Medicine, University of Cologne.

## FIGURE LEGENDS

**Supplemental Figure 1.**
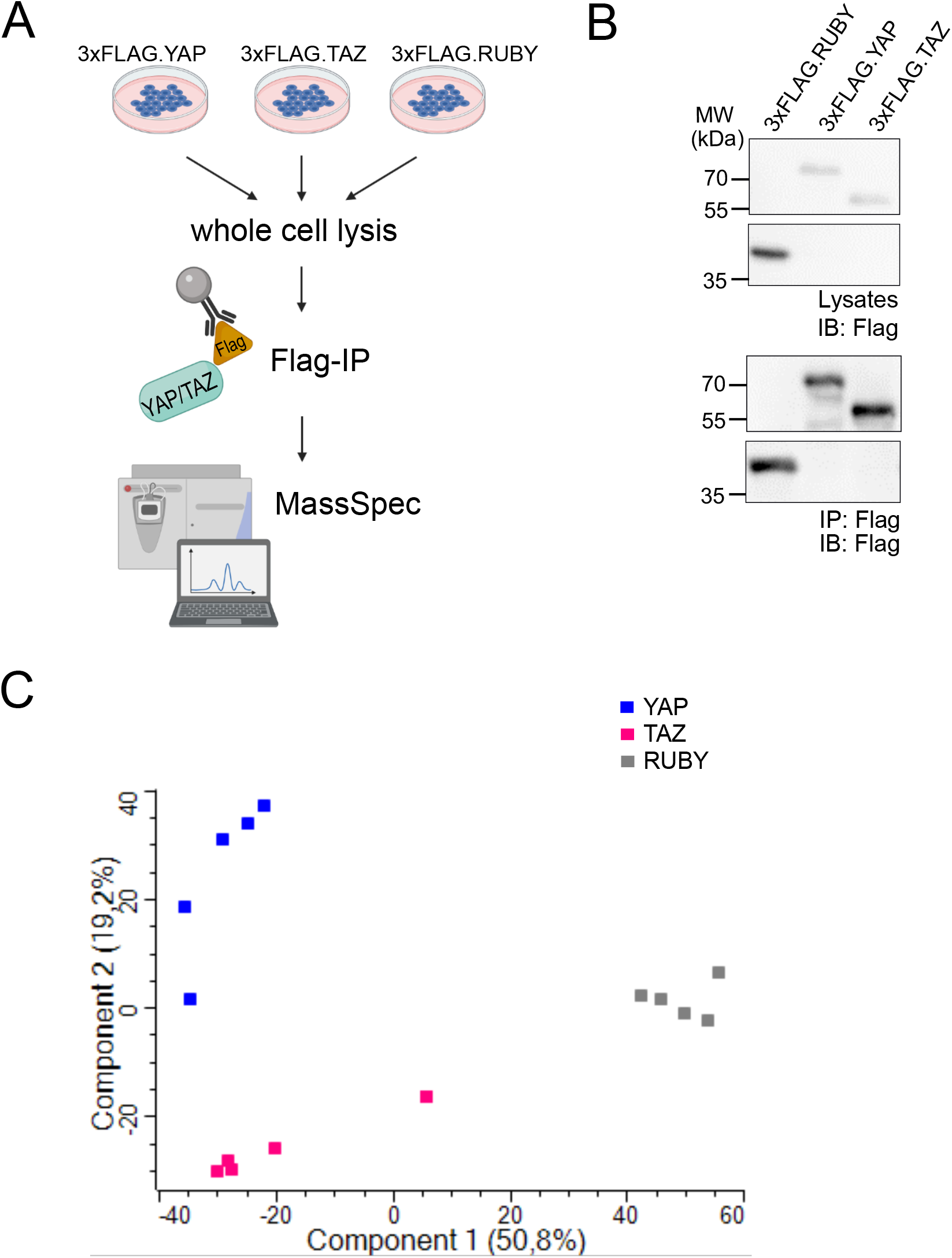
Mass Spectrometry sample submission and quality controls of the interaction datasets. A) Flag-immunoprecipitations of whole cells lysates of hsMPs cell lines overexpressing either flagged Ruby (3XFLAG.RUBY), Yap (3XFLAG.YAP) or Taz (3XFLAG.TAZ) were submitted to Mass Spectrometry (n=5). Created with BioRender.com. B) For every sample preparation an immunoblot was performed, showing successful pull-down of 3XFLAG.RUBY, 3XFLAG.YAP and 3XFLAG.TAZ. Representative data from the independent experiments is shown. C) Principal component analysis of all samples submitted. Scatter plot calculated from the label-free quantification (LFQ) values of all proteins identified in the respective samples: 3XFLAG.YAP in blue (YAP), 3XFLAG.TAZ in pink (TAZ) and 3XFLAG.RUBY in grey (RUBY).

**Supplemental Figure 2.**
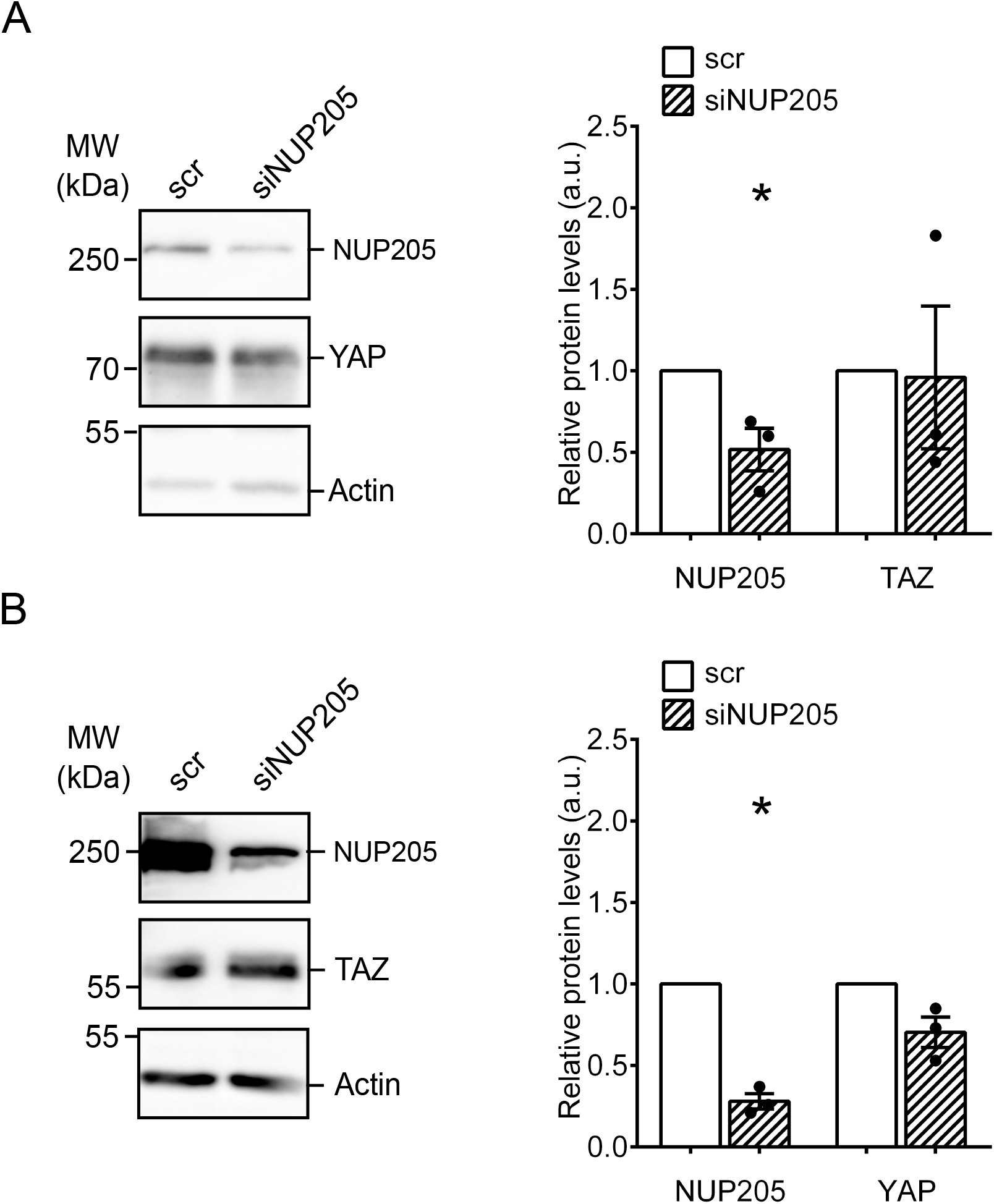
Downregulation of endogenous NUP205 does not affect YAP and TAZ whole cell endogenous expression. A) Dowregulation of endogenous NUP205 in HEK293T did not change YAP endogenous whole cells expression. B) Dowregulation of endogenous NUP205 in HEK293T did not change TAZ endogenous whole cells expression. Densitometry analysis of expression of YAP and TAZ relative to ß-Actin (arbitrary units,au). n=3. Mean ± SEM. * significant difference when compared to scrambled (p < 0.05; unpaired Student’s *t* test).

**Supplemental Figure 3.**
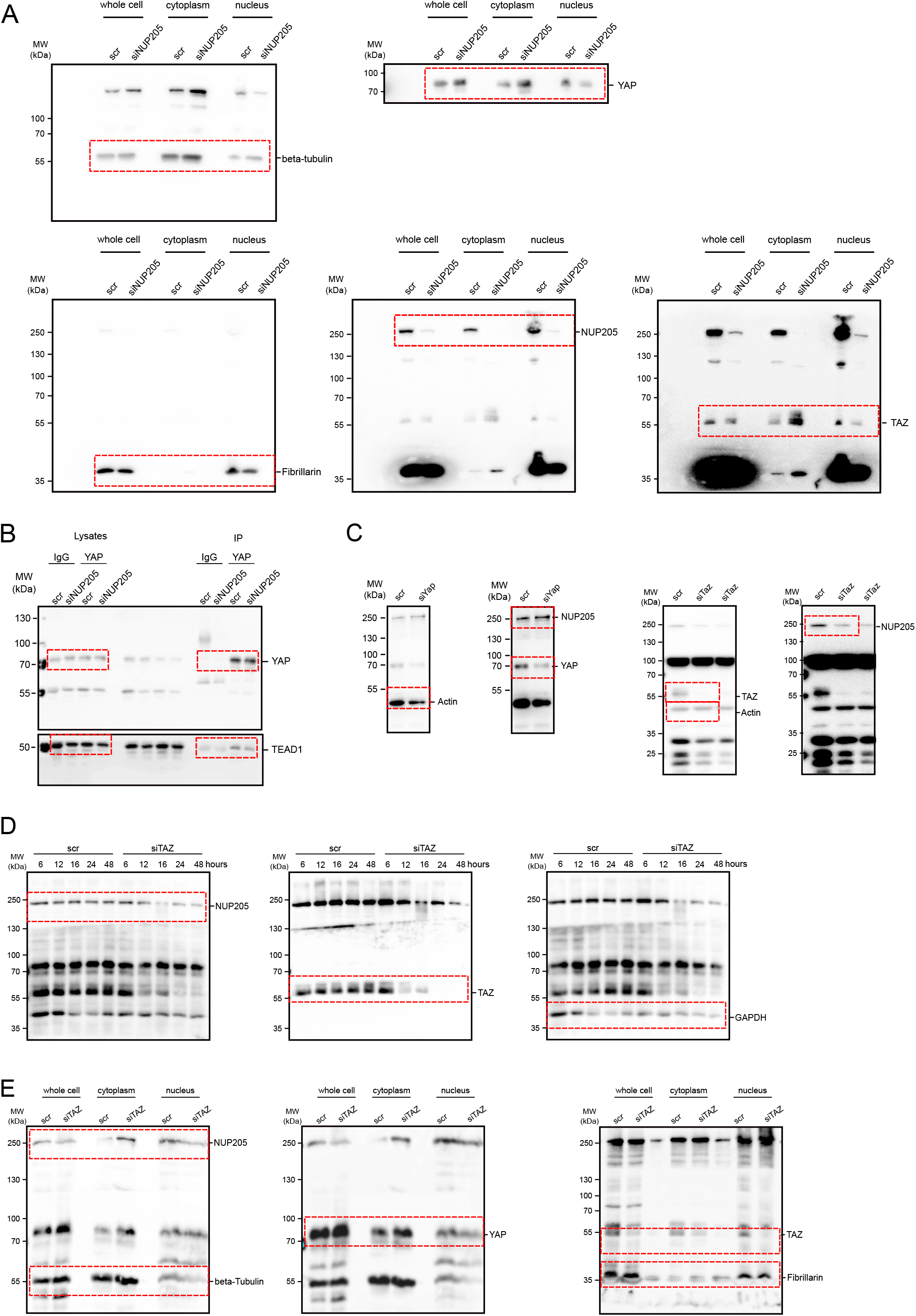
Original data: full-sized immunoblots. Original western blots, only cropped to membrane size (A) of Figure 2A, (B) of Figure 3A, (C) of Figure 4A, (D) of Figure 4B and (E) of Figure 4C.

## Notes

### Competing Interest Statement

The authors have declared no competing interest.

